# Tau-induced increase in promoter-proximal RNA polymerase II pausing is linked to suppressed expression of long neuronal genes in a *Drosophila* tauopathy model

**DOI:** 10.64898/2026.03.28.709859

**Authors:** Kendall Cottingham, Nima Goodarzi, David Fries, Geethma Lirushie, Hana Hall

**Affiliations:** Biochemistry Department, Purdue University, West Lafayette, Indiana, USA; Purdue Institute for Integrative Neuroscience, Purdue University, West Lafayette, Indiana, USA; Purdue Institute for Cancer Research, Purdue University, West Lafayette, Indiana, USA

**Keywords:** aging, taupathy, *Drosophila*, neurodegeneration, Alzheimer’s disease, promoter-proximal pausing, gene expression

## Abstract

Tauopathies, including Alzheimer’s disease, are age-related neurodegenerative disorders characterized by abnormal phosphorylation and buildup of microtubule-associated protein tau. Gene expression dysregulation is a key molecular feature of tauopathies, but how aging and disease interact to disrupt crucial transcriptional regulators and pathways remains largely unknown. Here, we examined how pathological tau affects gene expression programs in age-related neurodegenerative disease using a well-established *Drosophila melanogaster* tauopathy model with neuronal expression of the toxic human tau^R406W^. Transcriptomic analysis of tau-expressing fly heads showed a preferential downregulation of long neuronal genes with long introns. Notably, we found that these downregulated genes in the tauopathy model are marked by increased accumulation of initiating RNA polymerase II (RNAP II) near the transcription start site and reduced elongating RNAP II within gene bodies, indicating a problem with the transition from initiation to elongation. By calculating an RNAP II Pause Index (PI) for each gene, we identified a strong link between promoter-proximal RNAP II stalling, gene expression deficits, and gene length in the tauopathy model. Overall, we have uncovered the genomic and transcriptomic features of tau-dependent downregulated genes and identified increased RNAP II promoter-proximal stalling as a significant mechanism of transcription stress in tauopathy.

## 1. Introduction

Tauopathies are a diverse group of age-related neurodegenerative disorders characterized by the abnormal accumulation of tau, a microtubule-associated protein essential for microtubule assembly and stability (Black et al., 1996; Ramírez et al., 1999). The presence of intracellular neurofibrillary tangles composed of hyperphosphorylated tau is a key pathological feature of Alzheimer’s disease (AD) and frontotemporal dementia with parkinsonism linked to chromosome 17 (FTDP-17) (Hutton, 2000). Mutations in the microtubule-associated protein tau (MAPT) gene that affect microtubule binding, phosphorylation, or aggregation are responsible for most FTDP-17 cases (Goedert and Spillantini, 2000). FTDP-17 primarily affects the frontal and temporal lobes of the brain, leading to progressive changes in behavior and personality, communication difficulties, and impairments in memory and motor skills.

Although tau is involved in the development of various neurodegenerative diseases, the cellular mechanisms that cause neuronal dysfunction and death are still not well understood. Understanding the pathways that promote tauopathy progression, especially in the context of aging—which is the main risk factor for the disease—is crucial for deepening our knowledge of the molecular basis of tau-driven neurodegeneration (Niccoli and Partridge, 2012). Gene expression is increasingly recognized as a key factor in the development of tauopathies in both mouse models and human patients (Miller et al., 2008, 2010; Tan et al., 2021). During aging, existing transcriptional programs undergo widespread changes, driven by factors such as accumulated DNA damage and epigenetic modifications (López-Otín et al., 2023). These changes are thought to cause many of the physical signs of aging and age-related diseases. Previously, we showed that long, highly expressed genes encoding proteins with neuronal functions exhibit reduced expression in aging neurons (Hall et al., 2017). Later studies have also supported a connection between gene structure, transcription levels, and cellular function across different tissues and organisms, in aging and neurodegenerative disease contexts (Barbash and Sakmar, 2017; Lopes et al., 2021; Jauregui-Lozano et al., 2022; Stoeger et al., 2022; Debès et al., 2023; Gyenis et al., 2023). However, how aging and disease interact to alter the cellular environment and affect key transcriptional regulators that control gene expression remains largely unknown.

To investigate how tau affects gene expression programs in age-related neurodegenerative disease, we used a well-established *Drosophila melanogaster* tauopathy model with neuronal expression of human tau carrying the R406W mutation. Using steady-state RNA sequencing (RNA-seq), we characterized the age- and tau-dependent transcriptome in tauopathy flies. In tau transgenic flies, we observed a preferential downregulation of long genes characterized by long introns that encode factors essential for neuronal structure and function. Additionally, using the RNA polymerase II (RNAP II)-targeted Cleavage Under Targets and Release Using Nuclease (CUT&RUN) approach, we analyzed tau-dependent transcription dynamics. Downregulation of long genes was strongly linked to increased RNAP II stalling near the transcription start site (TSS) and decreased RNAP II occupancy along the gene body in the tauopathy model, suggesting that impaired promoter-proximal pause release is a likely mechanism behind their reduced expression. Overall, our results reveal the transcriptional dynamics and genomic features of genes dysregulated in tauopathy and their relationship to progressive, age-related neurodegeneration. Importantly, promoter-proximal RNAP II stalling emerged as a key characteristic of long, downregulated neuronal genes, implicating tau-dependent dysregulation of the transition from transcription initiation to productive elongation as a possible molecular driver of transcriptional stress in tauopathy.

## 2. Materials and Methods

### 2.1 *Drosophila* Stocks and Maintenance

The following strains used in this study were from Bloomington *Drosophila* Stock Center: elav>Gal4 (BDSC #8765), UAS-lacZ (BDSC #8529), UAS-tau (BDSC #78851). The UAS-TAU^R406W^ strain was a gift from Mel Feany (Wittmann et al., 2001). Flies were maintained under a 12:12-hour light: dark cycle at 25°C on standard fly food. For aging experiments, flies were collected within three days post-eclosion and transferred to fresh food every two days.

### 2.2 Optic Neutralization

Optic neutralization was performed as previously described (Stegeman et al., 2018). Briefly, 10 male flies per genotype were aged to the designated time point, anesthetized on a CO_2_ pad, and mounted to glass slides using clear nail polish as adhesive. Live rhabdomeres were imaged using brightfield light microscopy at 20x magnification. For each genotype and time point, at least 70 rhabdomeres were assessed. Retinal degeneration was quantified in a blinded manner and expressed as the percentage of lost rhabdomeres relative to the total number of rhabdomeres.

### 2.3 Climbing Assay

Climbing assays were performed using groups of 25 male flies per genotype. Flies were sorted into vials and allowed to recover for 24 hours following anesthesia. The next day, flies were gently tapped into a 100 ml glass graduated cylinder and acclimated for 2 minutes. The cylinder was then tapped firmly and swiftly three times to bring the flies to the bottom. Flies were given 10 seconds to climb, and the number of flies above a 5 cm mark was recorded. Each biological replicate consisted of three technical trials, with a one-minute rest between tests. For each genotype, 6-12 biological replicates (a total number of 150-300 flies) were assessed.

### 2.4 Western Blotting

Western blot analysis was performed using 40 μg of whole-cell protein extract from 25 fly heads per genotype. Heads were manually homogenized in cell lysis buffer (50 mM Tris-HCl, pH 8.0, 150mM NaCl, and 1% NP-40). Antibodies used for detection are listed in Supplemental Data 2.

### 2.5 RNA Extraction, Reverse Transcription-quantitative PCR (qPCR), and RNA-seq

Total RNA was extracted from 20-25 fly heads per genotype by mechanical homogenization in TRIzol reagent (Thermo Fisher, #15596018), followed by phenol/chloroform extraction and ethanol precipitation using standard protocols. cDNA was synthesized from 300-400 ng of total RNA using Episcript RNaseH Reverse Transcriptase (Biosearch Technologies, #ERT12925K-ENZ) with random hexamer primers (IDT). Quantitative PCR (qPCR) was performed using PR1MA qMAX Green 2X qPCR mix (Midsci, #PR2000-N-5000) on a CFX Connect Real-Time System (BioRad) according to the manufacturer’s instructions. Relative gene expression was normalized to the geometric mean of two reference genes in accordance with MIQE guidelines (Bustin et al., 2009). Primer sequences are provided in Supplemental Data 2.

For RNA-seq, extracted RNA was treated with DNase I (Roche, #04716728001) to remove genomic DNA according to the manufacturer’s protocol. Purified RNA was quantified using the Qubit RNA HS Assay Kit (ThermoFisher, #Q32852). cDNA libraries were prepared from 250 ng of RNA using the Universal Plus™ Total RNA-Seq Library Preparation Kit, incorporating *Drosophila*-specific anyDeplete technology for rRNA depletion (Tecan, #9156). Libraries were pooled and sequenced using paired-end 150 bp reads in a single lane on an Illumina Novaseq X Plus platform.

### 2.6 Bioinformatic Analysis of RNA-seq Data

Bioinformatic analysis of RNA sequencing data was performed as previously described (Jauregui-Lozano et al., 2022) with minor adjustments. Raw reads were trimmed using Trimmomatic (Bolger et al., 2014) to remove adapter sequences and low-quality bases. Cleaned reads were then quantified using Salmon (Patro et al., 2017) against a custom reference transcriptome built from *Drosophila melanogaster* (BDGP6.54) cDNA sequences, with the addition of *Homo sapiens* MAPT transcript information extracted from the GRCh38 cDNA reference. The resulting transcript-level counts were summarized to gene level and analyzed in R using the DESeq2 (Love et al., 2014) package for differential expression analysis. Cut-offs for differential gene expression were adjusted to a p-value < 0.05 and |fold change| > 1.5 unless otherwise stated. For comparisons between differential expression groups, statistical analysis was performed using ANOVA followed by a Tukey HSD post-hoc test. Gene Ontology (GO) enrichment analysis of significantly differentially expressed genes was performed using clusterProfiler (Yu et al., 2012).

### 2.7 CUT&RUN

CUT&RUN was performed using the EpiCypher CUTANA protocol with modifications for nuclei isolation from fly head tissue. Briefly, 100 fly heads were homogenized in 1 mL of Nuclear Extraction Buffer (20 mM HEPES, pH 7.9, 10 mM KCl, 0.1% Triton-X, 20% glycerol, 0.5 mM spermidine, 1× protease inhibitor) using a 1 mL Dounce homogenizer. Homogenization comprised 7 loose pestle strokes, a 5-minute ice incubation, followed by 7 loose and 12 tight pestle strokes. The homogenate was filtered through a 70 μm cell strainer, and 100 μL of nuclei was incubated with 11 μL of activated ConA beads per reaction. Targeted MNase cleavage was directed by antibodies listed in Supplemental Data 2. DNA was purified using the Zymo ChIP DNA Clean & Concentrator Kit (#D5205), and libraries were prepared with the EpiCypher CUTANA Library Preparation Kit (#14-1001). The libraries were pooled and sequenced in a single lane using paired-end 150-bp reads on an Illumina Novaseq X Plus platform.

### 2.8 Bioinformatics Analysis of CUT&RUN Data

Raw paired-end reads were trimmed and filtered for adapter sequences and low-quality bases using Fastp (Chen et al., 2018). Cleaned reads were aligned to the *Drosophila melanogaster* BDGP6.46 reference genome (annotation release 113) using Bowtie2 (Langmead and Salzberg, 2012) with end-to-end, sensitive parameters. The resulting BAM files were sorted with SAMtools and duplicate reads removed with Picard MarkDuplicates. Peak calling was performed using MACS3 (Zhang et al., 2008) with broad peak settings. The resulting pileup bedGraph files were normalized to counts per million (CPM) based on the total number of mapped reads and converted to BigWig format using bedGraphToBigWig from the UCSC Genome Toolkit. Normalized BigWig files were used to visualize metagene and signal profiles with deepTools (Ramírez et al., 2016), using the computeMatrix and plotProfile functions. For peak analysis, consensus peaks (defined as peaks present in at least three out of four biological replicates per condition) were identified using bedtools multiinter (Quinlan and Hall, 2010).

The RNAP II pause index (PI) is a gene-specific quantitative measure that assesses tau-dependent changes in promoter-proximal pausing relative to controls. Specifically, the Ser5-P pause index was calculated as the ratio of promoter Ser5-P RNAP II signal (transcription start site [TSS] ± 250 bp) to gene body Ser5-P RNAP II signal (+500 bp from the TSS to −500 bp from the transcription end site [TES]). Given the defined coordinates for promoter and gene body regions, a minimum gene length cutoff of 1500 bp was applied. Total Ser2-P was calculated independently as the average elongating RNAP II signal across the defined gene body region, serving as a direct measure of productive elongation. To determine tau-dependent differences in pause status, log2 fold changes of both the Ser5-P pause index and total Ser2-P signal were computed between tau-expressing and control flies on a per-gene basis. Overall, the RNAP II PI can capture relative changes in promoter-proximal RNAP II accumulation and relative levels of the elongating RNAP II signal in the tauopathy model. RNAP II PI can be mathematically defined as:

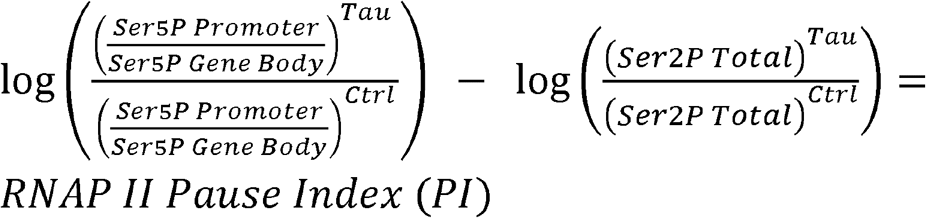

## 3. Results

### 3.1 Tau^R406W^ induces progressive, age-related neurodegeneration in a *Drosophila* tauopathy model

The tau R406W mutation, linked to FTDP-17 (Hutton, 2000; Nakamura et al., 2019), occurs at a conserved residue of exon 13 within the C-terminal domain, affecting all tau isoforms (Figure 1A) (Hong et al., 1998). Importantly, this mutant form of tau has structural and biochemical properties similar to those found in AD and FTDP-17 patients (Hutton, 2000). Tau R406W promotes hyperphosphorylation of tau, impairs its ability to bind microtubules, and increases neuronal vulnerability to apoptotic stress, thereby contributing to neuronal cell loss observed in tauopathy (Krishnamurthy and Johnson, 2004). In mouse models, tau R406W causes tau hyperphosphorylation and filament aggregation, disrupting axonal transport in affected neurons (Tatebayashi et al., 2002; Zhang et al., 2004; Ikeda et al., 2005). In *Drosophila*, pan-neuronal expression of transgenic human tau R406W (elav>htau^R406W^) (Figure S1A-C) reproduces key features of human tauopathies, including DNA damage (Frost et al., 2014), impaired locomotion (Merlo et al., 2014), and neuronal cell death (Wittmann et al., 2001). Using optic neutralization of the fly eye, we demonstrated that pan-neuronal expression of toxic htau R406W results in significant photoreceptor neuron loss starting in young, 10-day-old flies, which markedly worsens by middle age, 20 days after eclosion (Figure 1B, C, Figure S1D). In contrast, neuronal expression of wild-type human tau (WT-htau) does not cause retinal degeneration even in older flies, 40 days post-eclosion (Figure S1E). Notably, no rhabdomere loss was observed in 2-day-old early adult htau^R406W^ transgenic flies, indicating that tau-induced neurodegeneration is age-dependent and not due to retinal developmental defects. Similarly, progressive adult-onset brain vacuolization, a well-established marker of neurodegeneration in flies, has also been reported in this model (Wittmann et al., 2001).

**Figure 1.**
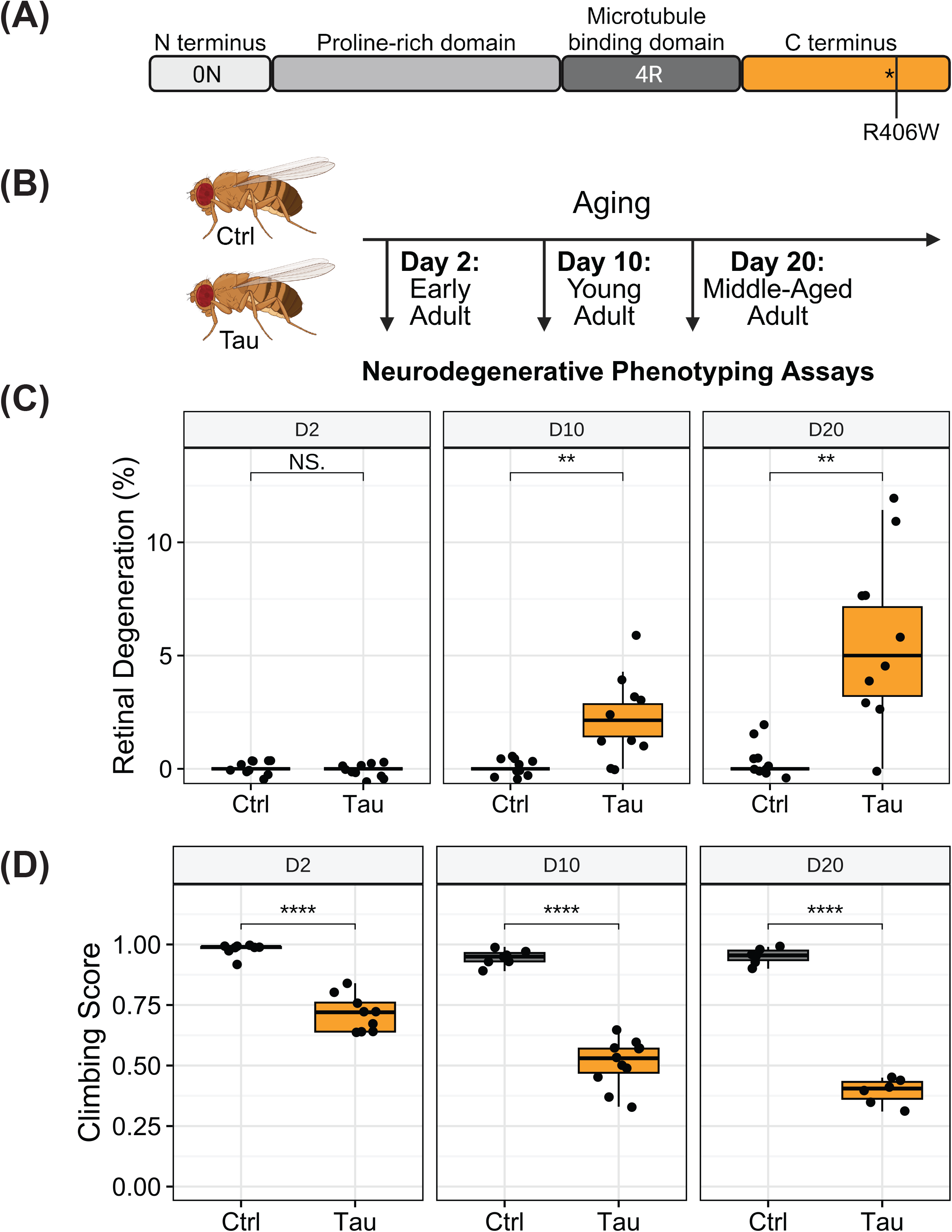
**(A)** Schematic of the human tau transgene isoform showing defined domains and the location of the R406W mutation. **(B)** Schematic illustrating the ageing time points used for phenotypic analysis. **(C)** Boxplot quantification of retinal degeneration in aged control (*elav>lacZ*) and tau (*elav>htau*^*R406W*^) flies. Rhabdomeres of 10 individual ommatidia per fly (n=10 male flies) were assessed. Retinal degeneration scores were calculated blindly as the ratio of the number of lost rhabdomeres to the total number of rhabdomeres, indicating photoreceptor cell death. **(D)** Boxplots showing the climbing ability of aged control (*Cyo/ htau*^*R406W*^) and tau (*elav>htau*^*R406W*^) flies. 6-12 replicates of 25 male flies were assessed for each genotype. Student’s t-test was used to determine statistical significance between genotypes at each time point (**** p < 0.001, **p < 0.01, *p< 0.05, NS/ns not significant).

To further assess age-related neurodegeneration in tau transgenic flies, we examined behavioral indicators of neuronal function. We carried out longitudinal climbing assays on htau^R406W^ transgenic flies and age-matched controls (no tau expression) to assess locomotion. Control flies maintained a high, consistent climbing score across different ages. In contrast, tau^R406W^ transgenic flies showed a 25% decrease in climbing ability at day 2 post-eclosion compared to controls (Figure 1D). This early-adulthood climbing-deficit phenotype suggests that htauR406W expression during development impairs neuronal function without affecting overall organization or survival. By day 10, htau^R406W^ transgenic flies demonstrated a 50% reduction in climbing ability relative to controls, with this decline progressing to a 70% reduction by day 20 (Figure 1D). The htau^R406W^ transgenic line, hereafter referred to as tau transgenic flies, exhibits quantifiable neurodegeneration early in life—before significant survival declines—making it an ideal model for studying mechanisms of tau-induced neuropathology (Wittmann et al., 2001). Additionally, they mimic the progressive, age-related neurodegenerative features seen in human tauopathies, highlighting their relevance as a *Drosophila* model of dementia (Creekmore et al., 2024).

**Supplementary Figure 1.**
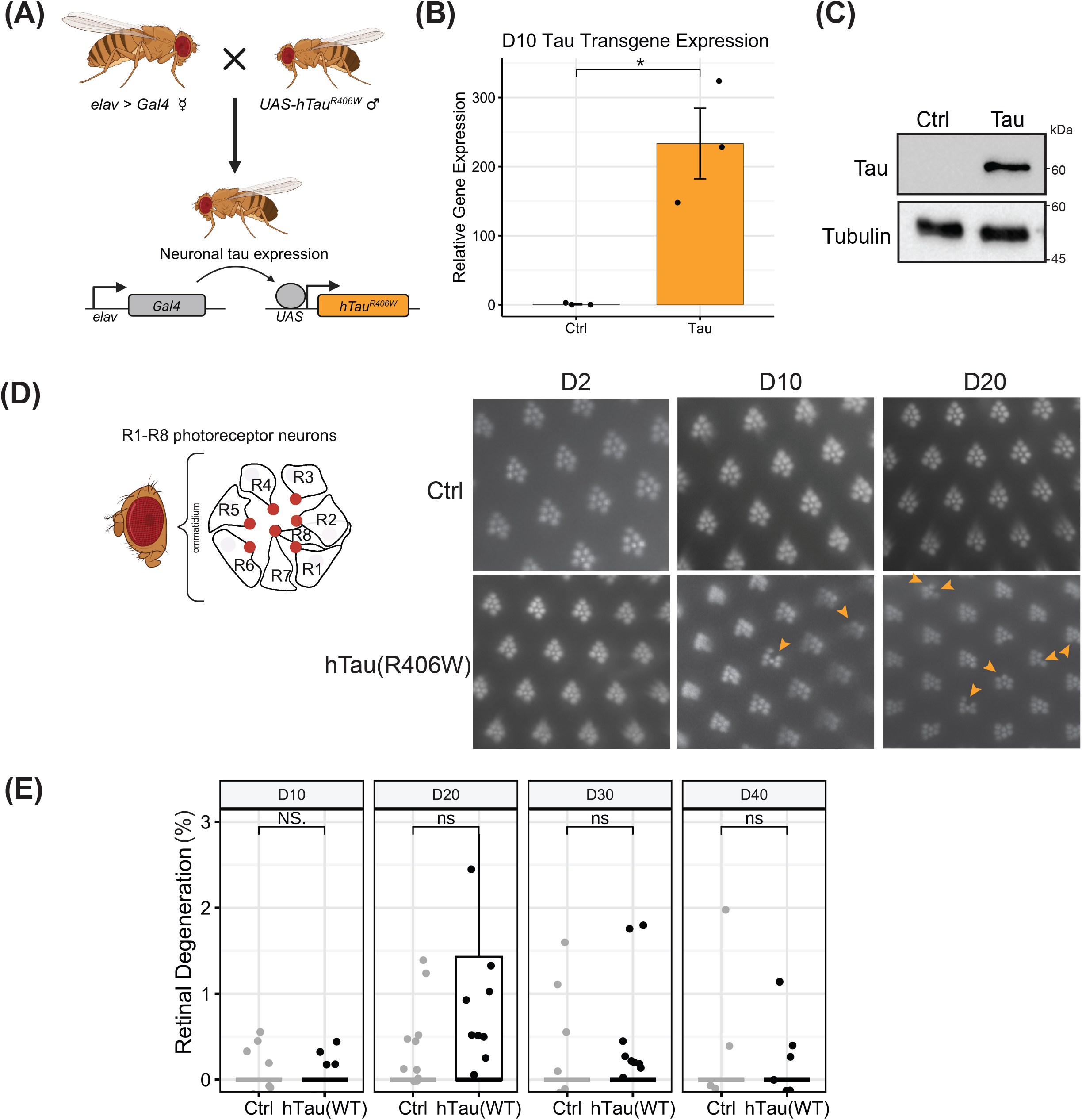
**(A)** A diagram illustrating a genetic approach to create a *Drosophila* tauopathy model using pan-neuronal *elav-Gal4*-driven expression of the toxic hTau^R406W^. Relative htau RNA **(B)** and protein **(C)** levels in 10-day-old transgenic flies. n=3 replicates of 25 male flies per genotype. **(D)** Left: Schematic of the *Drosophila* compound eye showing a single ommatidium enlarged to display photoreceptor neurons (R1-7) with rhabdomeres (red circles). Right: Representative images showing the loss of rhabdomeres in photoreceptor neurons (indicated by yellow arrowheads) in tau (*elav>htau*^*R406W*^) flies. **(E)** Boxplot quantification of retinal degeneration observed in aged flies expressing WT-htau. Student’s t-test was used to determine statistical significance between genotypes at each time point (NS/ns not significant).

### 3.2 Toxic tau expression drives dysregulation of neuronal, immune response, and cell cycle genes

Accumulation of tau in *Drosophila* and mouse models, as well as human brain tissue, leads to global chromatin relaxation and altered gene expression profiles, contributing to the development of neurodegenerative symptoms (Frost et al., 2014). Comprehensive transcriptomic analyses of human post-mortem brain tissue from individuals with AD and related tauopathies reveal distinct gene expression changes that correlate with cognitive and physical impairments (Mostafavi et al., 2018; Wan et al., 2020). Similarly, transcriptome profiling across various tau transgenic mouse models has revealed consistent transcriptomic alterations, including decreased expression of genes encoding synaptic proteins and increased expression of immune-response genes (Gjoneska et al., 2015; Kim et al., 2019). Thus, we sought to determine whether the tau-associated neurodegeneration observed in our *Drosophila* tauopathy model was accompanied by significant and clinically relevant alterations in canonical gene expression.

We conducted longitudinal RNA-seq using RNA extracted from the heads of control and tau transgenic flies at 2, 10, and 20 days post-eclosion (Supplemental Data 1). Three independent biological replicates, each comprising 20 male flies, were analyzed for each genotype and time point, except for a single discordant day 2 tauopathy fly sample, which was excluded from further analysis (Figure S2A and S2B). Principal component analysis (PCA) of the normalized transcript counts revealed that age accounted for 51% of the total variance across all samples, while tau expression contributed 18% (Figure S2A). Overall, the greatest variability in gene expression was observed among day 2 samples, likely reflecting dynamic transcriptional changes associated with the transition to adulthood. In contrast, samples from 10- and 20-day-old flies showed distinct genotype-specific clustering, indicating that tau expression is the primary driver of transcriptomic differences at these later stages (Figure S2B).

Longitudinal transcriptomic analysis revealed the greatest number of perturbations at day 10, with 1,640 differentially expressed genes (DEGs) identified in tau-expressing flies compared to controls (Figure 2A). In contrast, 704 and 805 DEGs were identified at days 2 and 20, respectively (Figure 2B 2C). Of all DEGs, 364 were consistently dysregulated across all three age-time points (Figure 2D), and these were enriched for gene ontology (GO) terms related to synaptic transmission and nervous system processes (Supplemental Data 1). This group includes 9 of the 10 genes encoding members of the synaptic nicotinic acetylcholine receptor (*nAChR*), a ligand-gated ion channel that regulates neuronal membrane potential. Decreases in human homologues of acetylcholine transporters and receptors yield cholinergic neuronal dysfunction in aging and AD (Nyakas et al., 2011). Consistently downregulated genes also included *Syt1* and *Tomosyn*, whose human orthologs, SYT1 and STXBP5, are significantly downregulated in postmortem AD brain tissue (Davidsson and Blennow, 1998; Canchi et al., 2019). The consistent, age-independent dysregulation of this gene subset suggests that toxic tau expression intrinsically alters the expression of key neuronal genes.

**Figure 2.**
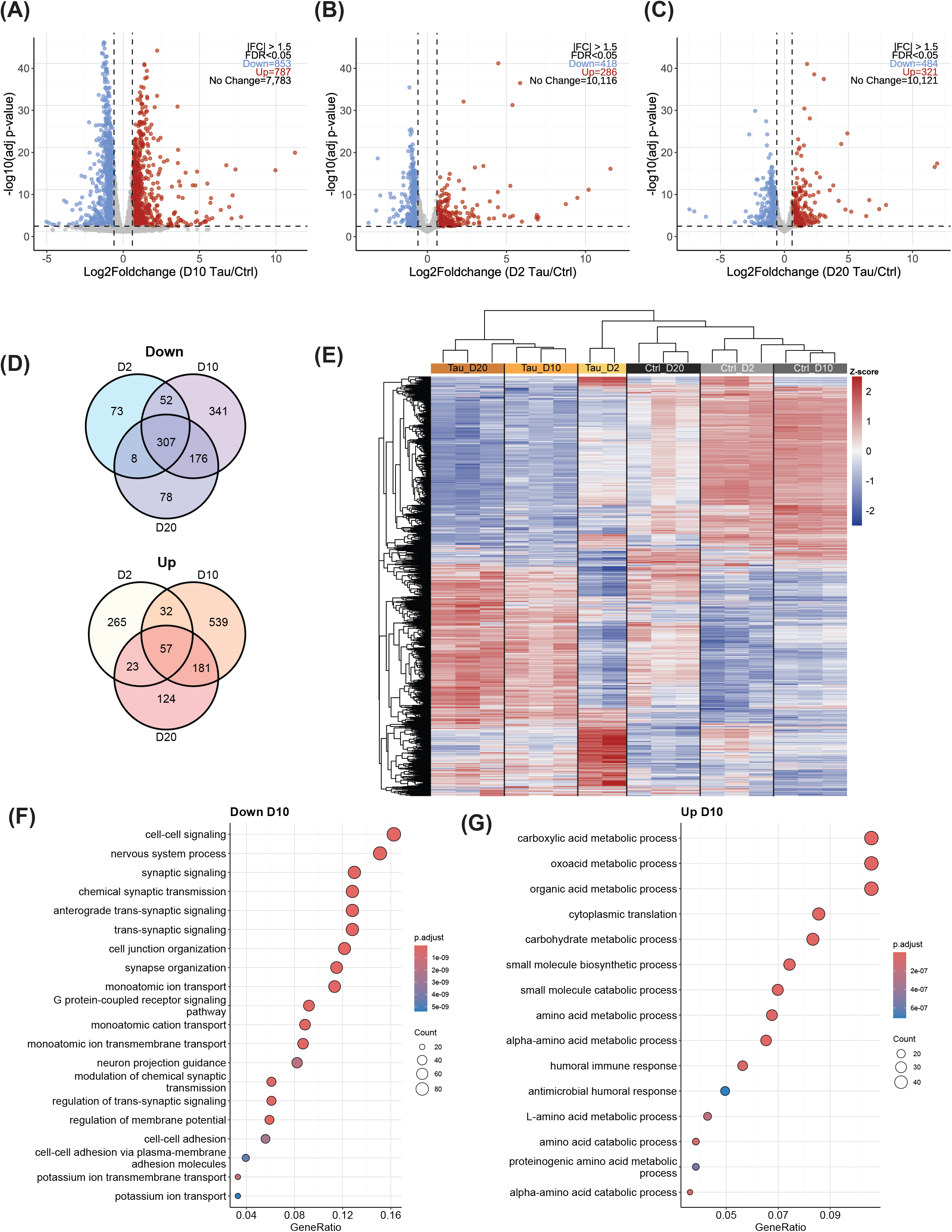
**(A)** Volcano plot showing differentially expressed genes (DEGs) between tau and control flies at day 2, **(B)** day 10, and **(C)** day 20, post-eclosion. DEGs are identified using DESeq2 (adjusted p value < 0.05, |FC| > 1.5). **(D)** Venn diagram illustrating the overlap between downregulated and upregulated genes across all aging timepoints. **(E)** Heatmap showing trends in expression Z-scores for all genes across all samples and aging timepoints. **(F)** Dot plot displaying the 20 most significantly enriched biological process gene ontology (GO) terms for downregulated genes in day 10 tau transgenics. **(G)** Dot plot illustrating the 15 most significantly enriched biological process GO terms for upregulated genes in day 10 tau transgenics.

Notably, gene expression progressively changed with age in 10- and 20-day-old control flies (Figure 2E). Importantly, the genes showing age-dependent expression changes in controls exhibited exacerbated dysregulation in tauopathy flies, emerging as early as 2 days of age and intensified markedly with age. These findings indicate that age-associated transcriptional shifts are accelerated in a pathological tau-dependent manner, highlighting the aging cellular environment as a key driver of tau-induced pathology and underscoring the profound impact of age on disease-associated expression signatures.

Given the extensive neuronal cell death observed in 20-day-old tau transgenic flies, we focused our subsequent transcriptomic analysis on 10-day-old flies to reduce the confounding effects of neuronal cell loss on gene expression profiles (Figure 1B and Wittmann et al., 2001). At this age time point, 17.4% of all expressed genes were differentially expressed in tau transgenic flies, with 853 genes being downregulated and 787 genes upregulated (p-adj < 0.05, |FC| > 1.5) (Figure 2B, Supplemental Data). We validated the RNA-seq findings using quantitative PCR (qPCR) on independent replicates of 10-day-old control and tau transgenic flies, targeting 10 genes identified as significantly down- or up-regulated in the transcriptomic dataset (Figure S2C, S2D).

Consistent with previous transcriptomic studies in tau transgenic models (Gjoneska et al., 2015; Mangleburg et al., 2020), GO term analysis of downregulated genes revealed enrichment for categories related to synaptic organization and neuronal function, including nervous system processes, chemical synaptic transmission, and regulation of membrane potential (Figure 2F). Among these, *hiw*, a gene encoding a RING E3 ubiquitin ligase playing an evolutionarily conserved role in axonal development, was notably downregulated. Its human ortholog, *MYCBP2*, has been linked to autism-like neurological symptoms when mutated (AlAbdi et al., 2023). Another gene, *Ten-a*, was also downregulated in 10-day-old tau-expressing flies, and, notably, a potential AD-related signal has been observed near the human TENM3 ortholog, associated with amyloidosis and tau pathology in AD cohorts (Deming et al., 2018). Additionally, *Gad1*, which encodes the enzyme responsible for synthesizing the inhibitory neurotransmitter gamma-aminobutyric acid (GABA), was downregulated in our tauopathy model. This mirrors the findings in chemically inactivated neurons, the aging human frontal cortex, and the hippocampus of AD patients, suggesting that reduced neuronal inhibition is a shared feature of aging and AD in both flies and humans (Blalock et al., 2004; Lu et al., 2004; Gleichmann et al., 2012).

In 10-day-old tau transgenic flies, upregulated genes were significantly enriched for GO terms associated with anabolic biological processes, such as small molecule biosynthesis and cytoplasmic translation, as well as various catabolic pathways, including amino acid and carbohydrate metabolism (Figure 2G). Notably, several cytochrome P450 genes - *Cyt-b5-r, Cyp4d1, Cyp4e2, Cyp9b1*, and *Cyp6a17* - were upregulated, aligning with human studies that report elevated levels of cytochrome P450 components in the cerebrospinal fluid and plasma in antemortem AD patients over healthy controls (Borkowski et al., 2021). Additionally, genes encoding proteins involved in immune response, particularly those from the immune deficiency (imd) and Toll signaling pathways, were markedly upregulated in tau transgenic flies. This robust activation of immune-related genes underscores chronic inflammation as a conserved feature of tauopathy in both flies and humans (Langworth-Green et al., 2023).

An increased body of evidence suggests that tau contributes to neurodegeneration by aberrantly reactivating the cell cycle in post-mitotic neurons, ultimately leading to apoptosis (Khurana et al., 2006; Frost et al., 2014; Zhao et al., 2021a). Supporting this mechanism, our transcriptomic analysis revealed upregulation of genes encoding proteins involved in DNA replication and cell cycle progression, including proliferating cell nuclear antigen (*PCNA*), *Nnf1b, Trithorax*-*like* (*Trl*), *Origin recognition complex subunit 1* (*Orc1*), and Minichromosome maintenance 2 (*Mcm2*). We also observed increased expression of *Nucleophosmin* (*Nph*), a gene that typically protects neurons but, when overexpressed, can promote cell cycle re-entry and apoptosis (Pfister and D’Mello, 2015). Conversely, gene expression of Wee1 kinase (*Wee1*), a key regulator of the G2/M checkpoint, was reduced, further supporting dysregulation of cell cycle control (Esposito et al., 2021).

Collectively, these findings highlight the similarities between gene expression changes in the *Drosophila* tauopathy model and those observed in human Alzheimer’s disease. They also help disentangle the distinct and overlapping contributions of aging and tau pathology to transcriptomic alterations. Importantly, the gene expression changes identified in the 10-day-old tau transgenic flies reveal early transcriptional disruptions associated with tau toxicity. These include reduced synaptic signaling, increased immune response, and aberrant cell cycle activation, all of which are key features of progressive tau-related neurodegeneration.

**Supplementary Figure 2.**
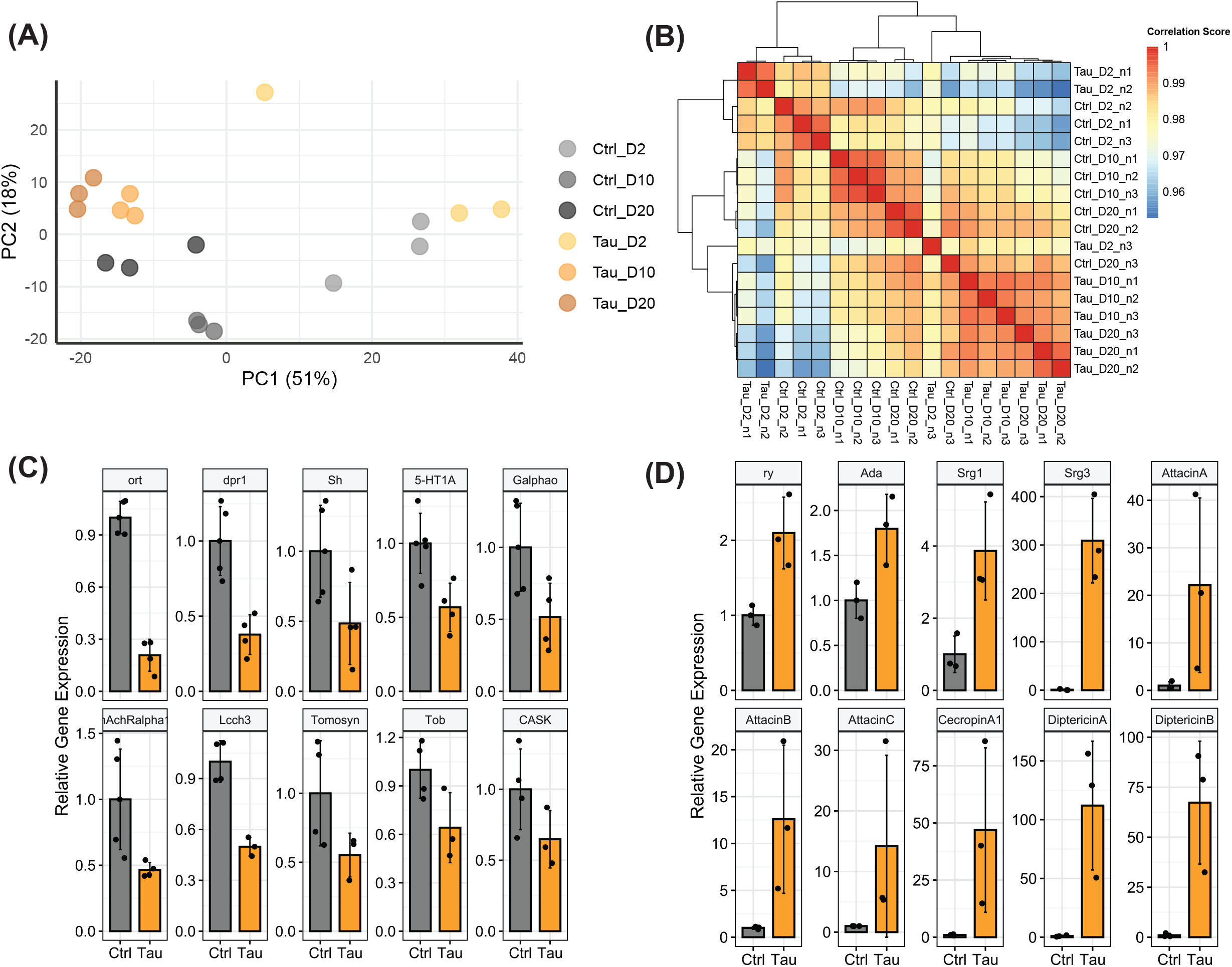
**(A)** Principal component analysis (PCA) of longitudinal RNA-sequencing samples based on normalized counts per million. **(B)** Hierarchically clustered heatmap of Pearson correlation coefficients calculated from normalized RNA-seq expression values across all samples. Bar plots displaying the relative transcript levels of genes assessed by qPCR that were identified as downregulated **(C)** or upregulated **(D)** in tau transgenic flies by RNA sequencing. n=3-4 replicates of 20 male flies per genotype.

### 3.3 Long genes with long introns are preferentially and significantly downregulated in tau transgenic *Drosophila*

We previously demonstrated that long, highly expressed genes encoding proteins with neuronal function tend to decrease expression in aging *Drosophila* photoreceptor neurons; a decline that correlates with age-related visual impairment (Hall et al., 2017). Similar associations between gene length, transcriptional output, and cellular function have been reported across various tissues and organisms in both aging and neurodegenerative disease contexts (Barbash and Sakmar, 2017; Lopes et al., 2021; Stoeger et al., 2022; Debès et al., 2023; Gyenis et al., 2023). To explore whether similar features characterize genes dysregulated in our tau transgenic flies, we analyzed transcriptomic data based on conserved genomic attributes and expression levels. In 10-day-old tau transgenic flies, downregulated genes were significantly longer than upregulated or unchanged genes (Figure 3A). Importantly, average transcript abundance did not differ significantly between downregulated genes and other differential expression groups (Figure S3A). GO term analysis of long downregulated genes (>10kb) revealed enrichment for categories associated with neuronal structure and function, including synapse organization and signaling (Figure 3B). Since neuronal genes are generally longer than those expressed in other cell types (Zylka et al., 2015), we controlled for potential bias by focusing on genes associated with GO0050877 (Nervous System Process), a highly enriched and broadly encompassing neuronal GO term among downregulated genes (Figure 2C). Even within this subset, downregulated genes were significantly longer than genes in other differential expression categories (Figure 3C). In contrast, long genes that did not change expression in tau transgenic flies were associated with cell morphogenesis and developmental processes (Figure 3D).

**Figure 3.**
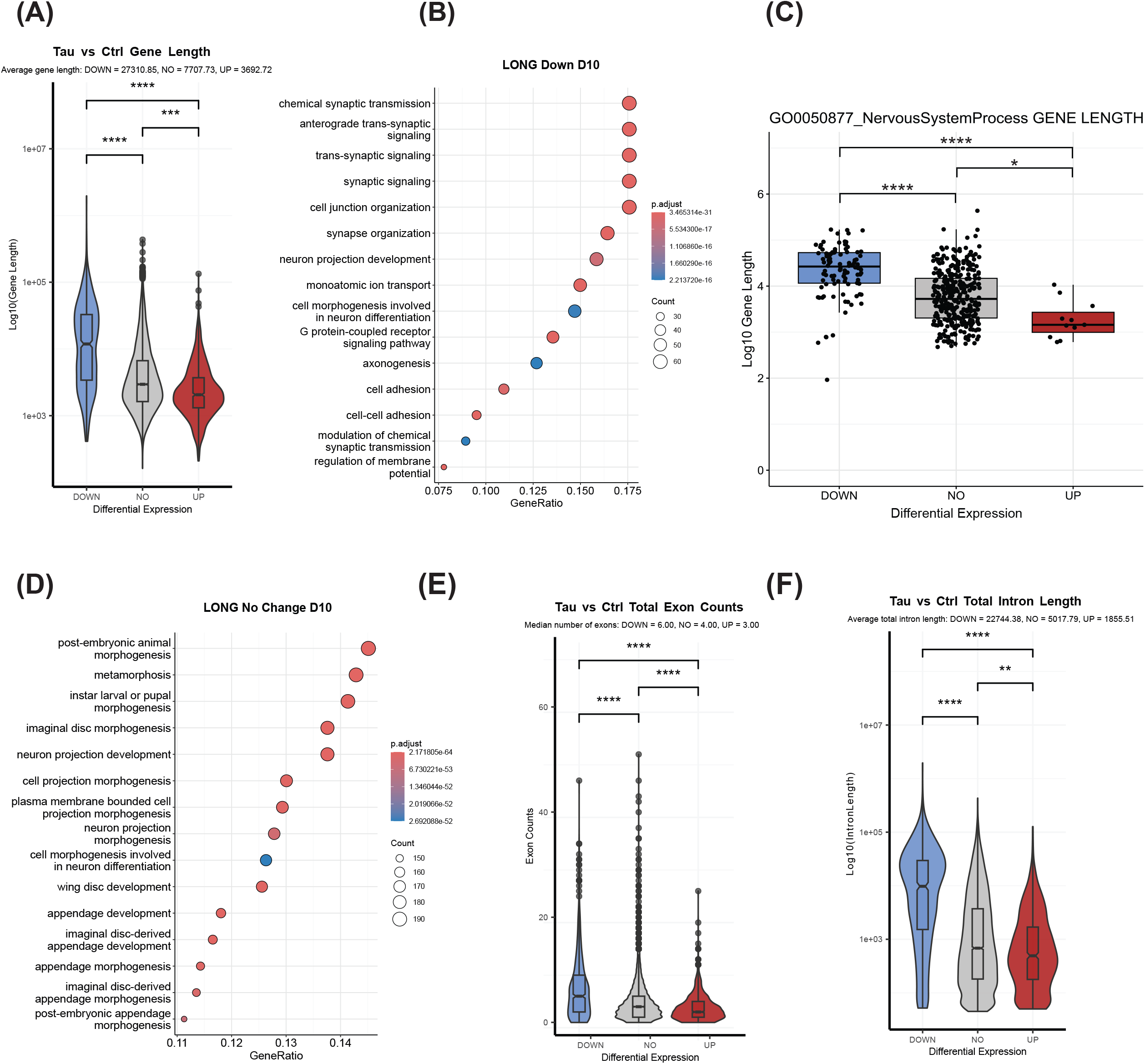
**(A)** Violin and boxplot of differentially expressed gene (DEG) categories showing the total length of all genes. **(B)** Dot plot displaying the 15 most enriched GO biological process terms associated with long, downregulated genes. **(C)** Boxplot showing the length of genes within ‘GO0050877: Nervous System Process’ by DEG categories. **(D)** Dot plot displaying the 15 most enriched GO biological process terms associated with long non-DEGs. Violin and boxplot of differentially expressed gene (DEG) categories showing **(E)** number of exons, and **(F)** total intron length. For differential expression data, DOWN (downregulated), NO (no change), and UP (upregulated) expression in tau transgenic flies compared to controls at day 10 post-eclosion. Long genes are defined as those > 10kb. ANOVA followed by TukeyHSD post-hoc test was used to evaluate statistical significance between DEG groups (**** p < 0.001, * p < 0.05, NS/ns not significant).

Gene length is shaped by both exon and intron architecture. While exons are short and highly conserved, introns vary widely in length and can impact transcription efficiency (Carmel et al., 2007; Yenerall and Zhou, 2012). In our tauopathy model, long genes that were significantly downregulated contained more exons than genes with unchanged expression and approximately twice as many as those in the upregulated gene group (Figure 3E). However, the total length of their introns was four times that of genes with unchanged expression, averaging approximately 22 kb, compared with approximately 5 kb for introns in genes with unchanged expression (Figure 3F). Long intron splicing deficits in the tauopathy model may contribute to gene dysregulation; however, at the transcript level, there is no change in the expression of key splicing factors (Figure S3B). Overall, the observed difference in intron length largely explains the increased total gene length observed in downregulated genes, suggesting that this genomic feature may play a critical role in regulating proper gene expression in tauopathy flies.

**Supplementary Figure 3.**
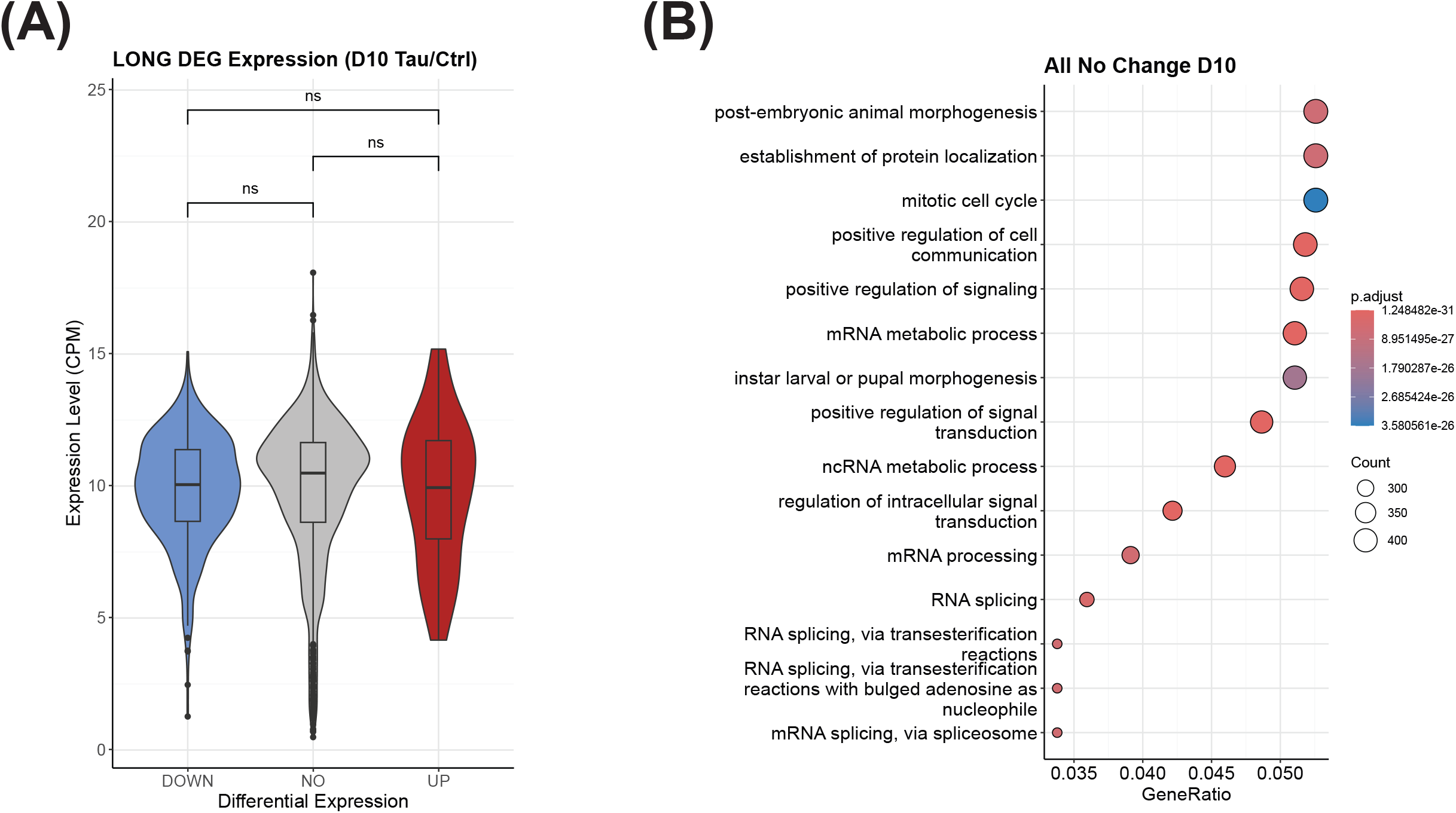
**(A)** Violin and boxplot illustrating the expression of all long genes based on DEG categories. Differential expression data are indicated as DOWN (downregulated), NO (no change), and UP (upregulated) expression in tau transgenic flies compared to controls at day 10 post-eclosion. Long genes are defined as > 10kb. ANOVA followed by TukeyHSD post-hoc test was used to determine statistical significance between DEG groups (**** p < 0.001, *** p < 0.005, ** p < 0.01, *p< 0.05, NS/ns, not significant). **(B)** Dot plot displaying the 15 most enriched GO terms among all non-DEGs.

### 3.4 Increased promoter-proximal RNA polymerase II pausing is associated with long downregulated genes

To focus our investigation of tau-dependent dysregulation of gene expression, we assessed transcription dynamics using genome-wide profiling of RNAP II. After transcription initiation, RNAP II typically pauses near the promoter, and its timely release into productive elongation is a key regulatory step in fine-tuning gene expression. To map the RNAP II distribution in tau transgenic flies, we assayed 10-day-old tau and age-matched control flies using the CUT&RUN technique. This approach, combined with antibodies specific for Ser5- and Ser2-phosphorylated (Ser5- and Ser2-P) RNAP II, markers of initiating and elongating forms of the polymerase, respectively (Bozukova et al., 2022), enabled us to assess RNAP II dynamics with high resolution.

CUT&RUN generated 3-6 million uniquely mapped fragments across four biological replicates per genotype and antibody. Principal component analysis revealed that the primary source of variance was RNAP II phosphorylation state (PC1) and tau expression (PC2) (Figure S4A). In control flies, metaplot analysis revealed a sharp RNAP II peak around the TSS, followed by a plateau across gene bodies and a decrease at the transcription end site (TES) (Figure 4A, first column, gray line), consistent with a canonical RNAP II pattern across actively transcribed genes (Bozukova et al., 2022). In contrast, tau transgenic flies showed elevated levels of initiating RNAP II at the TSS and reduced levels of elongating RNAP II around the promoter and throughout the gene body (Figure 4A, first column, yellow line). Stratifying RNAP II profiles by gene expression revealed that both up- and downregulated genes in tau transgenic flies exhibited increased initiating RNAP II occupancy at the TSS (Figure 4A, top row). For upregulated genes, elongating RNAP II levels were comparable between tau-transgenic and control flies, suggesting that increased transcript levels may result from enhanced transcription initiation or increased RNA stability (Figure 4D, bottom row). Conversely, downregulated genes exhibited a marked depletion of elongating RNAP II across gene bodies, consistent with reduced transcriptional output.

**Figure 4.**
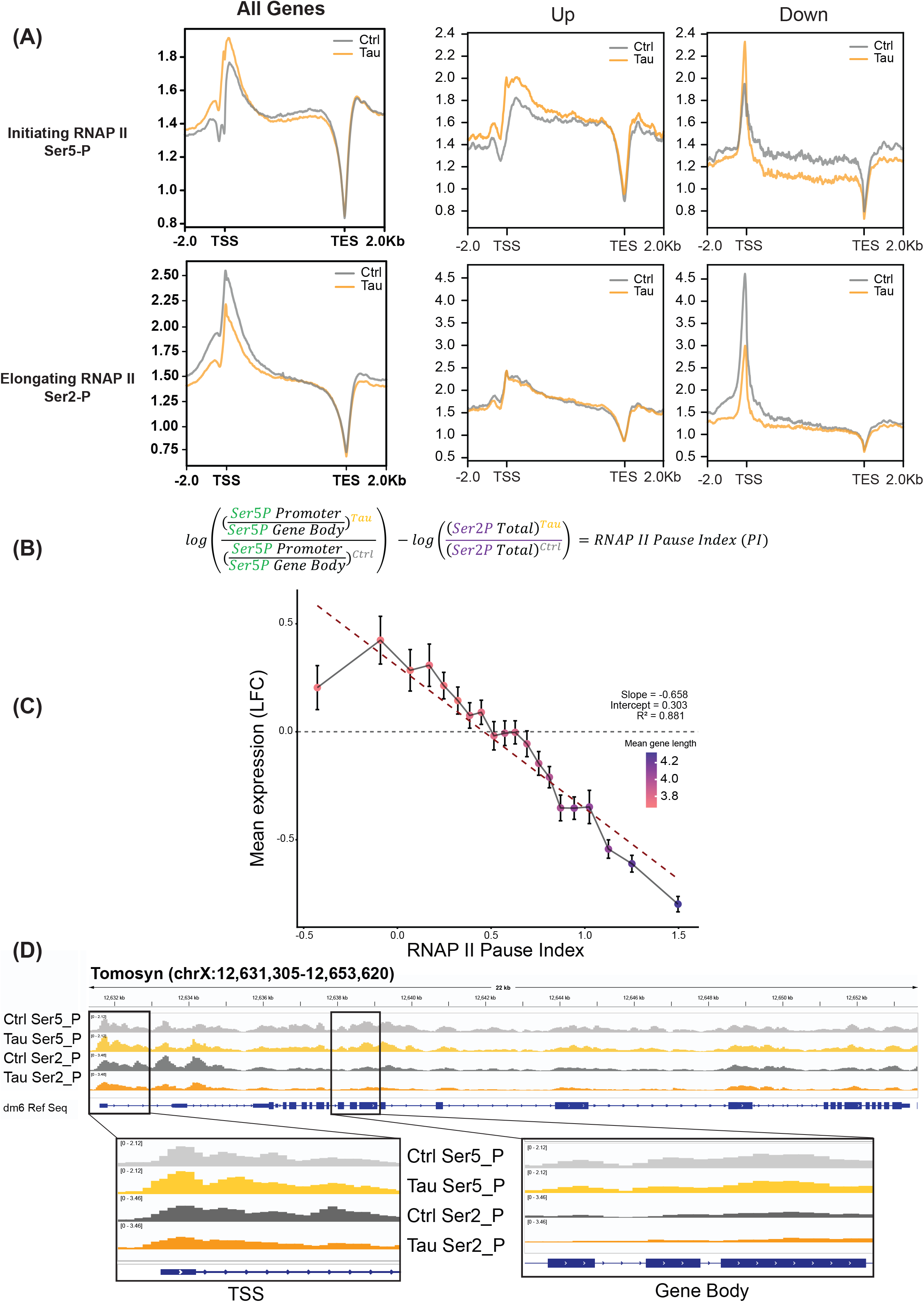
**(A)** Metaplots of CPM (counts per million)-normalized Ser2- and Ser5-phosphorylated RNAP II CUT&RUN signals over gene bodies of all expressed protein-coding genes, all up- and downregulated genes for control (gray) and tau transgenic flies (yellow). TSS, transcription start site; TES, transcription end site. **(B)** Calculations for determining RNAP II Pause Index (PI). PI values greater than 0 indicate increased promoter-proximal pausing for a given gene in the tauopathy model. **(C)** Dot plot showing RNAP II pausing index (PI) values, gene expression, and gene length. Genes were ranked by increasing PI score and grouped into 20 equal-sized bins. Bin-wise means of PI value, gene expression, and gene length were plotted to visualize global trends. **(D)** Genome browser inspection of RNAP II profiles in the Integrated Genomic Viewer (IGV) depicting a representative long, downregulated neuronal gene, *Tomosyn*, from the highest RNAP II PI bin. RNAP II density represents the average of four independent biological replicates per genotype and RNAP II phosphor-isoform.

To more specifically quantify pause proclivity in the tauopathy model, we calculated the RNAP II Pause Index (PI), an initiation-to-elongation transition factor, for each gene. Building on previously described RNAP II pause-release ratios (Akhtar et al., 2021), we integrated our phospho-specific RNAP II dataset by defining the PI as the ratio of promoter-enriched Ser5-P RNAP II to total gene body Ser2-P RNAP II occupancy for each gene in tauopathy versus control flies (Figure 4B). Using this RNAP II PI as an unbiased, gene-level measurement of promoter-proximal pausing, we uncovered a strong relationship between polymerase pausing, gene expression changes, and gene length (Figure 4C, Figure S4B). Specifically, genes exhibiting significantly increased promoter-proximal pausing were long and downregulated in tau-transgenic flies. Further, within the most-paused gene cohort, those with the highest average RNAP II PI, we identified several long, downregulated neuronal genes, including *Tomosyn* and *Frq1*, that exemplify the canonical pattern of increased promoter proximal RNAP II pausing (Figure 4D, E).

These findings reveal that RNAP II dynamics are specifically disrupted in the tauopathy model. Notably, downregulated genes in tau transgenic flies are marked by the concomitant accumulation of initiating RNAP II and depletion of elongating RNAP II. Taken together, these RNAP II profiles suggest that exacerbated promoter-proximal polymerase pausing is a hallmark of long, downregulated genes in the tauopathy model. This promoter-proximal polymerase stalling may underlie failure of critical transcriptional processes, particularly the transition from initiation to elongation, as a potential mechanism driving transcription stress in tauopathy.

**Supplementary Figure 4.**
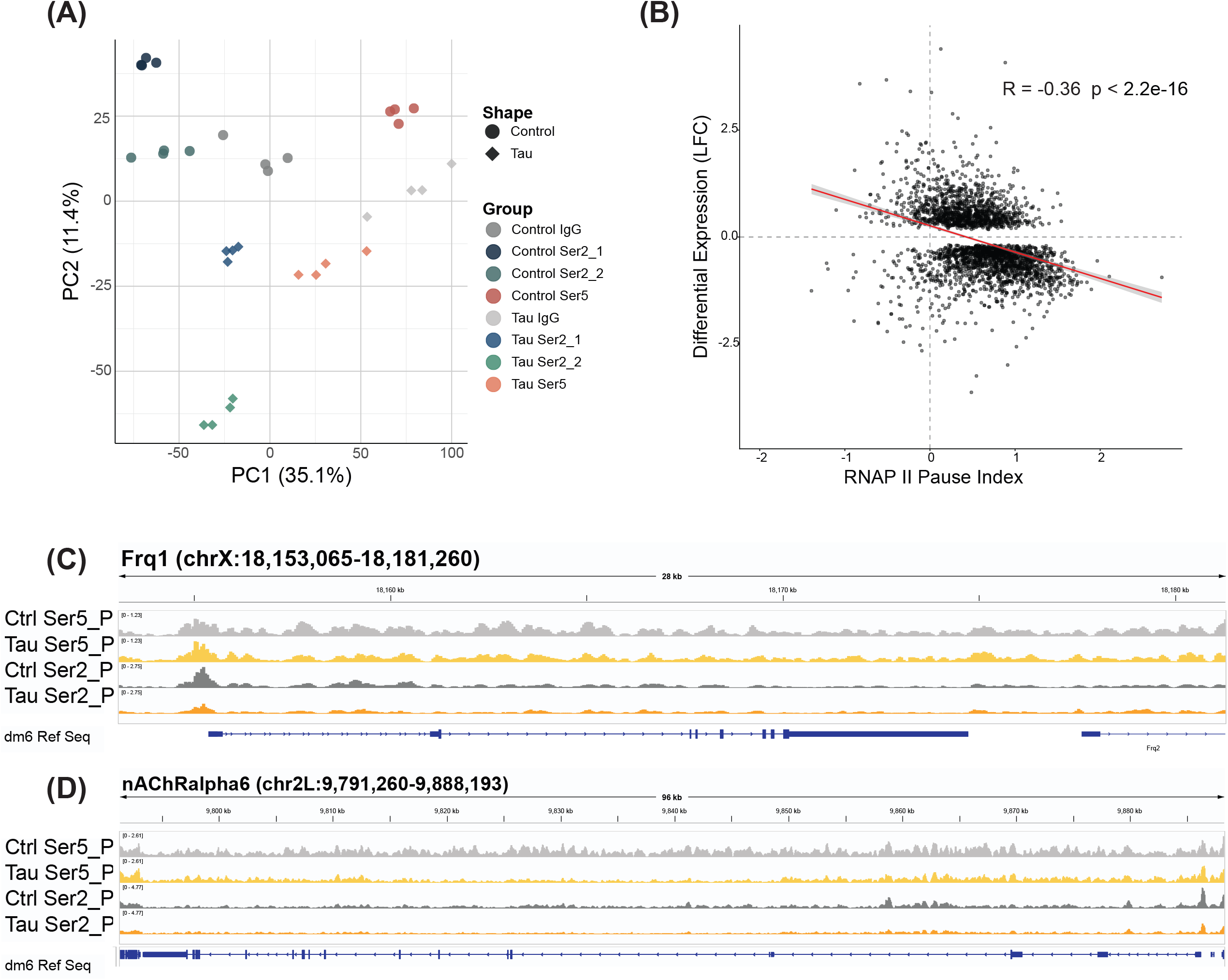
**(A)** PCA plot of RNAP II CUT&RUN samples based on consensus peak counts. The dot shape indicates genotype, and the dot color indicates antibody target. **(B)** Dot plot showing the correlation between RNAP II PI and gene expression for all expressed genes with an adjusted p-value <0.05. Genome browser inspection of RNAP II profiles in the Integrated Genomic Viewer (IGV) depicting representative long, downregulated neuronal genes, *Frq1 (C)* and *nAChRalpha6 (D)*, from the highest RNAP II PI bin. RNAP II density represents the average of four independent biological replicates per genotype and RNAP II phosphor-isoform.

## 4. Discussion

Gene dysregulation is a central hallmark of tauopathies, including Alzheimer’s disease, and underpins a range of pathological symptoms. Despite growing recognition of its importance, our understanding of the multilayered regulation of gene expression, encompassing transcriptional, epigenetic, and post-transcriptional mechanisms, in neurodegenerative disease remains limited, particularly in the context of aging (Miller et al., 2008, 2010; Wan et al., 2020; Tan et al., 2021). In this study, we used a well-established *Drosophila* tauopathy model with neuronal expression of toxic human tau^R406W^. Previously, the age-dependent transcriptome and proteome of this tauopathy model were characterized in a comparative analysis to identify intersectional trends in RNA and protein expression (Mangleburg et al., 2020). Here, we further interrogated the ubiquitous and age-dependent transcriptional alterations driven by tau accumulation and identified conserved genomic features of downregulated genes to elucidate the underlying molecular mechanism driving gene dysregulation.

Tau-expressing flies exhibited preferential downregulation of genes required for maintaining neuronal structure and function, including those involved in learning, memory, and locomotion. Concurrently, we observed upregulation of genes associated with stress and immune responses, metabolism, and cell cycle regulation. This tau-dependent gene-dysregulation profile recapitulates age-related changes in expression. However, tau markedly accelerates and amplifies age-dependent transcriptional stress, underscoring how aging and disease converge to shape pathological gene expression signatures. These disrupted gene expression patterns also mirror findings from other tau transgenic models, including aberrant cell cycle re-entry of post-mitotic neurons, a process widely implicated as a key driver of neurodegeneration in tauopathies and AD (Khurana et al., 2006; Zhao et al., 2021b).

Characterization of distinct genomic trends among differentially expressed genes in the tauopathy model revealed that downregulated genes were significantly longer and had a four-fold increase in cumulative intron length compared to non-differentially expressed genes. While neuronal transcripts are generally longer than those of other cell types (Zylka et al., 2015), the subset of downregulated neuronal genes in our dataset was notably longer than other neuron-specific genes that were not differentially expressed. These findings align with a growing body of research indicating an inverse relationship between gene length and transcriptional output in the context of aging and neurodegenerative disease (Zylka et al., 2015; Hall et al., 2017; Gyenis et al., 2023).

Further analysis revealed that long, downregulated genes in tau-expressing flies showed reduced elongating RNAP II occupancy across gene bodies. These downregulated genes also showed an accumulation of the initiating form of RNAP II just downstream of the transcription start site, suggesting a failure to release promoter-proximal paused polymerase into active elongation. On active genes, the transition of primed RNAP II into elongation is inhibited by negative regulators NELF and DSIF, promoted by P-TEFb-mediated phosphorylation, and maintained by other key histone modifiers and RNA-binding proteins (Jonkers and Lis, 2015). At the transcript level, we observed no significant changes in the expression of these critical elongation factors in the tauopathy fly model (data not shown). However, recent studies have shown that transcription factors, splicing factors, and RNA-binding proteins can co-localize with pathogenic tau tangles in the cytoplasm (Hsieh et al., 2019; Kavanagh et al., 2022), potentially limiting their nuclear availability and regulatory function.

Notably, recent work in *Drosophila* shows that the nuclear exon junction core complex (pre-EJC) associates with promoter regions through interactions with the initiating, Ser5-P RNAP II, mediated by nascent RNA transcripts, thereby reinforcing promoter-proximal pausing. This effect is particularly pronounced at genes containing long introns (Akhtar et al., 2019). In this context, our findings further underscore, in particular, gene and intron length as critical determinants of tau-induced transcriptional stress, suggesting that genes with these attributes are especially susceptible to transcriptional dysregulation.

Gene dysregulation is a defining molecular hallmark of age-related diseases; yet, how these regulatory pathways become disrupted in specific disease contexts remains a central question in neurodegenerative disease biology. Our data reveal distinct transcriptional dynamics and genomic features associated with downregulated transcripts in tauopathy, providing compelling evidence that promoter-proximal RNAP II stalling is a hallmark of long neuronal genes. These findings underscore the importance of studying gene regulatory mechanisms within disease-relevant contexts and highlight a pathological shift in transcriptional control in tauopathy. Future studies that elucidate the key factors in tau pathology that drive promoter-proximal RNAP II stalling and further clarify the connection between genomic features, such as gene and intron length, and RNAP II dynamics at vulnerable neuronal genes will help unravel the complexity of transcriptional regulation in age-related tauopathy. Furthermore, given the close interplay between genomic architecture, nascent RNA modifications, and chromatin structures in transcription regulation, investigating how these co-transcriptional elements collectively shape gene expression in neurodegeneration will be of significant interest.

## Author Contributions

KC: Data curation, Formal Analysis, Investigation, Methodology, Visualization, Writing – original draft, review, and editing. NG: Formal analysis, Visualization, Writing–review. DF: Validation, Writing–review. GL: Validation, Writing–review. HH: Conceptualization, Funding acquisition, Resources, Supervision, Writing-review, and editing.

## Data Availability Statement

The RNA-seq and RNAP II CUT&RUN datasets generated for this study are available in the GEO repository [link].

